# Designer DNA nanocages modulate anti-oxidative and anti-inflammatory responses in tumor associated macrophages

**DOI:** 10.1101/2024.07.01.601504

**Authors:** Payal Vaswani, Dhiraj Bhatia

## Abstract

Cancer is a complex disease, with multiple treatment modalities, but no definitive cure. The tumor microenvironment contributes to the complexity of the disease by forming a niche of multiple cell types supporting each other to carry out various cellular functions. Tumor associated macrophages are one such kind of cells which support the tumor microenvironment via immunosuppression. DNA tetrahedron (TD), a widely explored DNA nanocage, has shown a lot of potential in therapeutics. However, the role of TD still remains quite unexplored in immunology. Here, we first establish the anti-oxidative and anti-inflammatory role of TD. We then proceed with using TD as a therapeutic agent in tumor associated macrophages by modulating the response of PD-L1. The findings of this work create a base for TD in biological applications such as cancer immunotherapy.

**Graphical Abstract:** Immunomodulatory effects of DNA nanocages on tumour associated macrophages

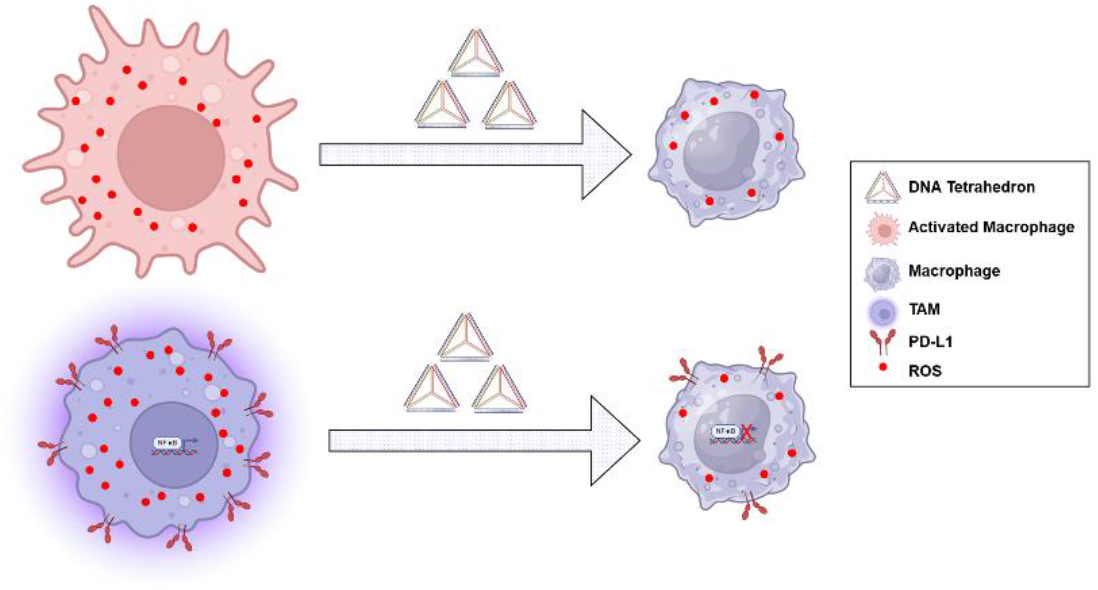

## 1. Introduction

Cancer is an age-old disease with still a lot to unfold. The primary treatment for cancer includes surgery, chemotherapy and radiotherapy. We are now in the transformative era of targeted therapies, immunotherapies and personalized medicine^1^. However, we are still struggling with finding a definitive cure. Cancer forms a complex tumor environment which includes cancer cells, stem cells, immune cells (macrophages, T cells), blood vessels, and extracellular matrix^2^. Tumor microenvironment can promote drug resistance and immune evasion which might present significant challenges in formulating effective treatment strategies^3^. Further advancements in cancer research are essential for deciphering the complexities and developing new and innovative treatments.

Macrophages are important part of our innate immune system^4^. They transition from resting stage (M0) to active stage (M1 and M2). M1 macrophages are generally termed as fighters as they defend against pathogens. On the other hand, M2 macrophages act as healers during tissue repair. In diseases such as cancer, macrophages are recruited as anti-cancer moieties. However, in the tumor microenvironment, the cancer cells repolarize the macrophage to pro-tumoral^5^. These macrophages are termed as tumor associated macrophages (TAM). They help the tumor in processes such as angiogenesis, tumor growth, metastasis, and creating an immunosuppressive environment^6^. The current therapies are focused on targeting the cancer cells but not on the entire tumor microenvironment^7^. Targeting TAMs alongside existing therapies could enhance overall improvement in cancer therapy.

DNA nanotechnology is a booming field focused on applications in disease treatment and therapeutic interventions. The principles of DNA nanotechnology utilize the Watson-Crick base pairing principles to design complex nanostructures^8^. Prof. Ned Seeman first formed 4-way junction marking the start of the field^9^. Since then, various structures such as DNA tetrahedron (TD)^10^, DNA icosahedron^11^, DNA cube^12^, and so on have emerged. TD is a nanocage with plethora of advantages and applications associated with it. It has been used for different applications such as drug delivery^13–15^, stem cell differentiation^16,17^, and gene silencing^18^ among others. TD has also shown anti-oxidative and anti-inflammatory activities^19^.

In this study, we explore the potential application of TD in tumor immunology. We synthesized TD and characterized it using various techniques. We first checked the anti-oxidative properties of TD followed by anti-inflammatory properties. We proceeded further by checking the effect of TD on the endocytosis mechanism in the M0 and activated macrophage cells. We established that TD alone does not cause any immune reaction. So, we further investigated the role of TD in TAM. Finally, we also explored the targets affected by TD in TAM. Our study will open up new avenues of TD in the field of immunology which is not yet fully explored.

## 2. Results

### 2.1 Synthesis and Characterization of Tetrahedron

We used one pot synthesis to carry out TD synthesis^20^ (**Figure 1a**). In brief, the four primers (M1, M2, M3 and M4) were mixed in equimolar ratio in 2 mM MgCl_2_ buffer, followed by thermal annealing. Size and morphology-based characterization of TD was carried out using EMSA, DLS and AFM. EMSA technique relies on band retardation based on molecular weight. We used 8% native PAGE to check the band pattern when different number of primers were added in the same ratio. We found that with increasing number of primers, the migration becomes slow, being the least in TD (**Figure 1b**). This shows higher order structure formation. We then moved to check the size of formed structure using DLS. DLS shows the hydrodynamic size based on light scattering. We found the hydrodynamic size of TD as 9.2 ± 0.78 nm (**Figure 1c**). We proceeded to check the morphology of the formed TD using AFM.

**Figure 1:**
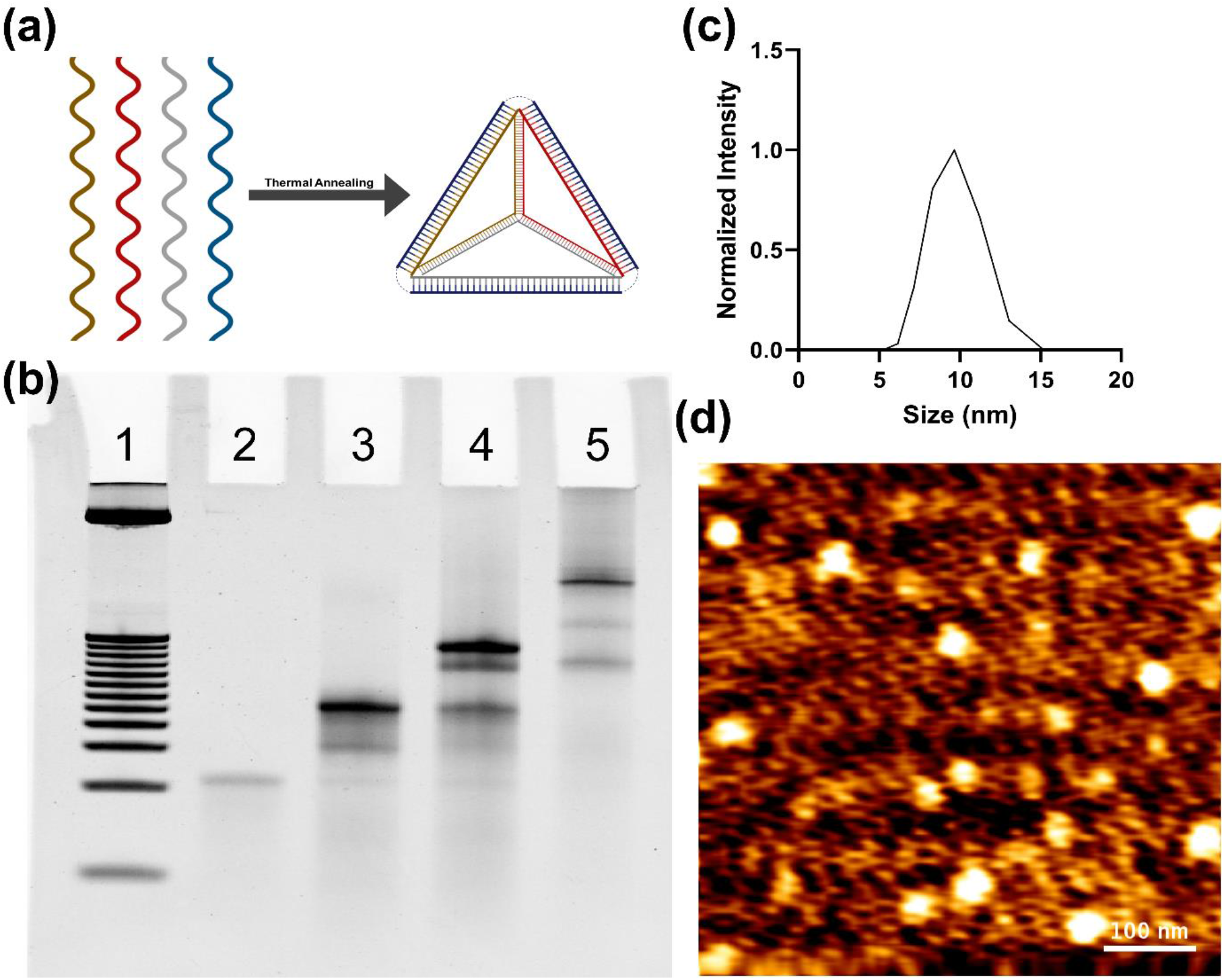
Synthesis and Characterization of DNA Tetrahedron (TD). **(a)** Schematic representation of TD synthesis. Four single stranded primers are subjected to thermal annealing to form TD **(b)** EMSA performed using 8% native PAGE. Lane 1: 25 bp DNA Ladder; Lane 2: M1; Lane 3: M1+M2; Lane 4: M1+M2+M3; Lane 5: M1+M2+M3+M4. Band retardation shows higher order structure formation. **(c)** DLS showing the hydrodynamic size of TD as 9.2 ± 0.78 nm. **(d)** AFM image showing the triangular morphology of TD.

We found triangular structure formation with size around 18-20 nm (**Figure 1d**). We finally carried out the stability assay for TD using serum and found it to be stable up to 2 hours after which it started degrading (**Supplementary Figure 1**).

### 2.2 Anti-oxidative and anti-inflammatory properties of tetrahedron

The levels of reactive oxygen species (ROS) are pivotal for the regulation of various cellular functions^21^. Increased ROS production is primary inflammatory response in macrophages. We first wanted to establish the effect of TD on ROS and to do that, we used RAW264.7 cells as our macrophage model. It is known that lipopolysaccharide (LPS) can polarize the macrophages to M1 state and in turn increase the ROS production^22^. So, we divided the cells into 5 groups: non-treated control, 200 nM TD for 2 hours, 500 ng/mL LPS for 24 hours, TD pretreatment followed by LPS treatment and post treatment of TD after the LPS treatment.

We used DCF-DA for checking the ROS. We found that the level of ROS in TD treated group is similar to our non-treated cells indicating that TD doesn’t have capability to induce ROS by itself (**Figure 2a, b**). LPS treated cells, which acted as our positive control, had the highest levels of ROS. The cells pre-treated and post-treated with TD had decreased levels of ROS compared to LPS. To verify the same, we repeated the same experiment using multimode plate reader which indicated the same pattern (**Supplementary Figure 2**). This suggests that TD does have anti-oxidative properties.

**Figure 2:**
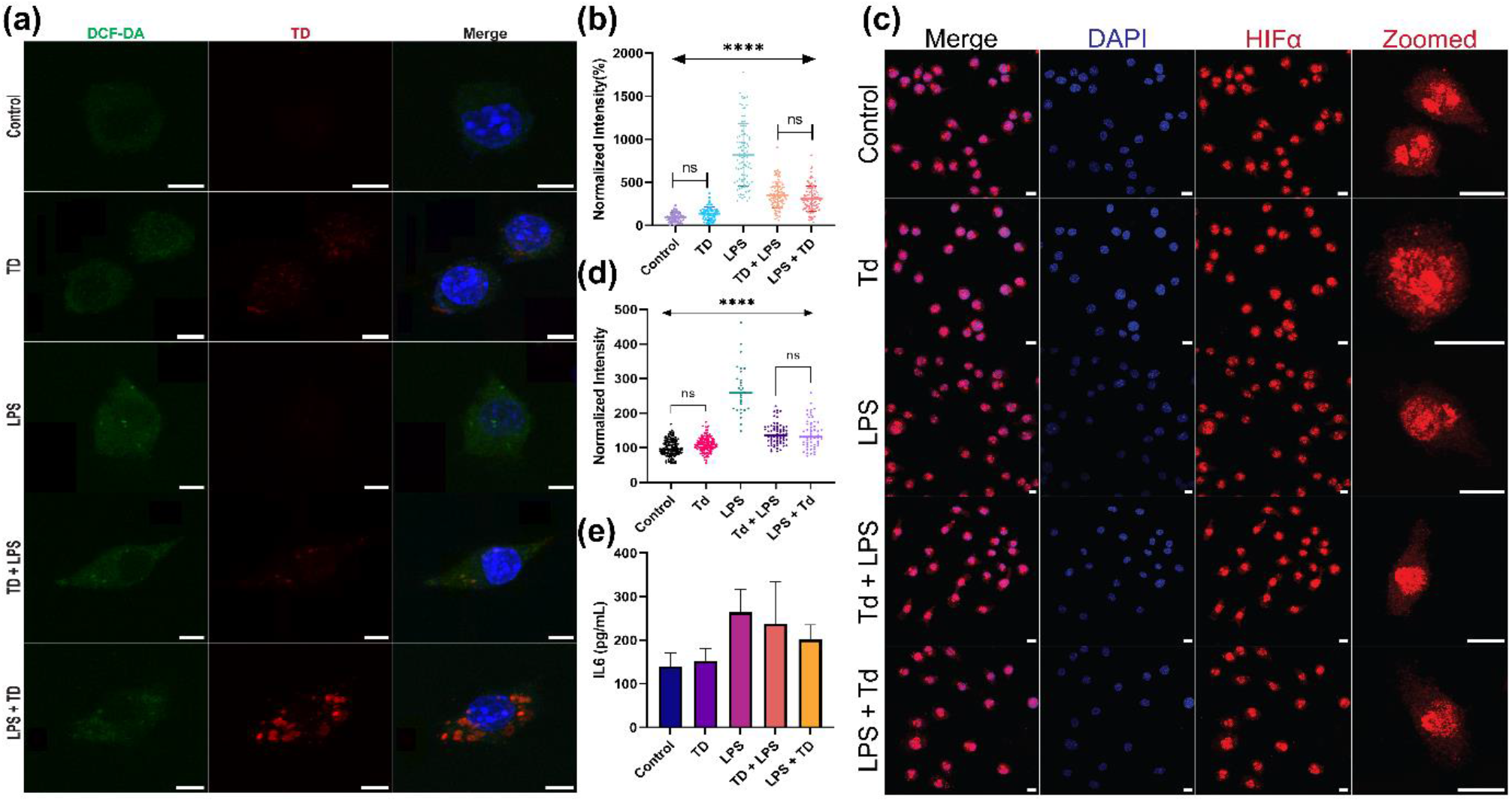
Anti-oxidative and anti-inflammatory properties of TD. **(a)** Confocal images of DCF-DA staining showing ROS levels in RAW264.7 cells. Green color indicated ROS levels by DCF-DA staining, red staining showing TDCy5 uptake and blue color indicated nucleus stained by DAPI. **(b)** Quantification of ROS levels of the images in panel (a). **(c)** Confocal images showing HIFα immunostaining. Blue color indicates DAPI staining of nucleus, red staining indicated HIFα. **(d)** Quantification of HIFα expression in the cells in panel (c). **(e)** IL6 levels in the five groups using ELISA. Scale bar is 20 µM.

Increased expression of HIFα has shown increased levels of ROS^23^. To check whether the similar correlation is applicable to macrophages, we carried out Hifα immunostaining. We found similar trend in the expression of HIFα to ROS levels, which validated our results (**Figure 2c, d**). ROS production is required for IL6 release in M1 macrophages^24^. IL6 is also one of the cytokines released by M1 macrophages^25^. So, we estimated the levels of IL6 in our treatment groups. We found that in cells treated with LPS, the IL6 levels were highest (**Figure 2e**). With pre- or post-treatment of TD, the levels of IL6 decreased compared to LPS. Hence, we showed that TD has anti-oxidative and anti-inflammatory properties.

### 2.3 Phagocytosis and Endocytosis

Macrophages are phagocytes and known for their ‘bacteria eating’ property^26^. To test the effect of our treatment on phagocytosis, we decided to carry out cellular uptake of FITC-dextran 40 kDa. As expected, the uptake of dextran was highest in the LPS condition (**Figure 3a, b**). With the treatment of TD, we observed that dextran uptake lowered suggesting decreased functionality of the macrophages.

**Figure 3:**
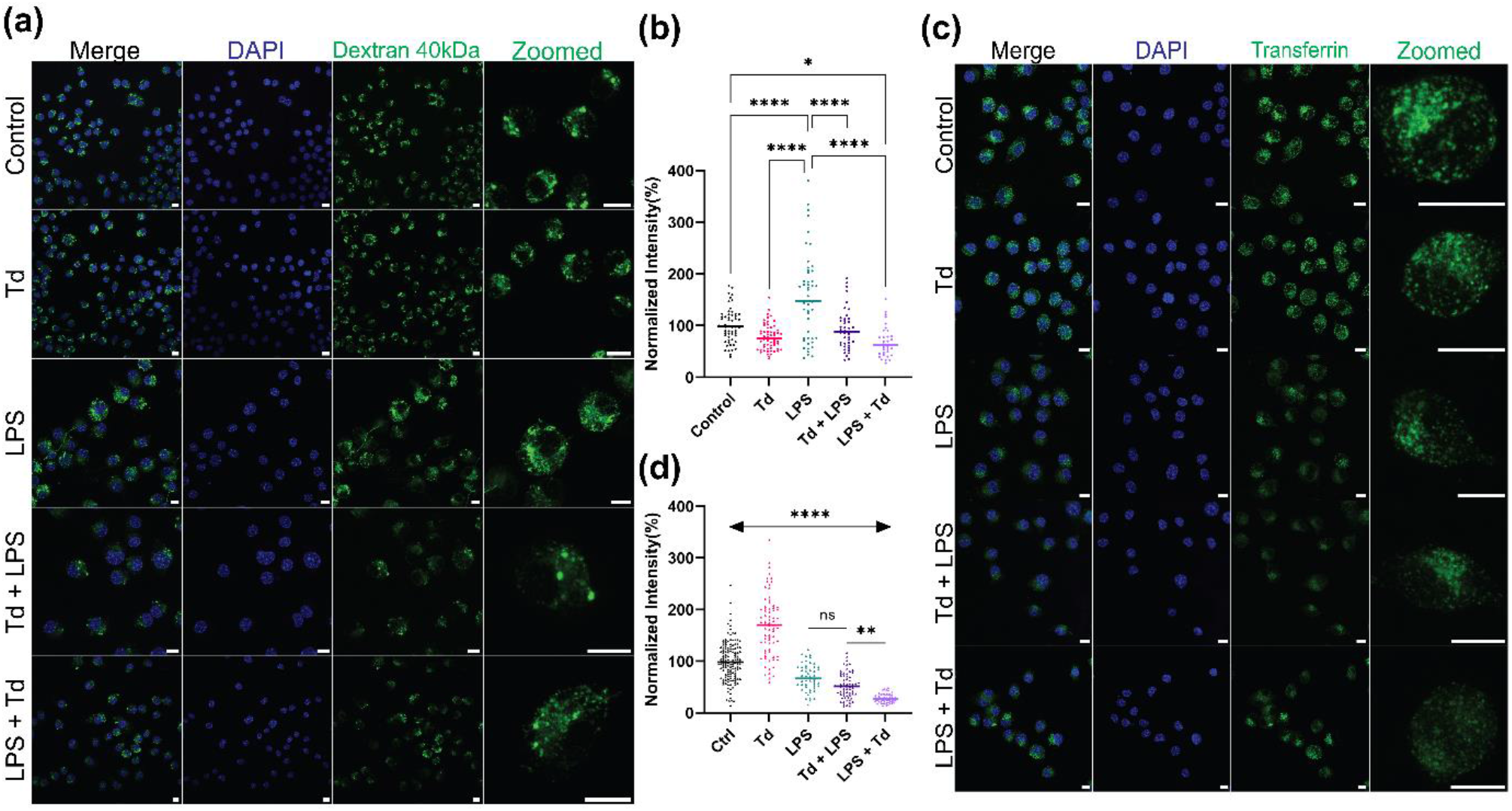
Phagocytosis and Endocytosis. **(a)** Confocal images showing cellular uptake of FITC dextran. *Blue color indicates DAPI staining of nucleus, green color indicates 40 kDa dextran*. **(b)** *Quantification of dextran uptake shown of images in panel (a)*. **(c)** Confocal images showing cellular uptake of transferrin-A647. *Blue color indicates DAPI staining of nucleus, green color indicates transferrin*. **(d)** *Quantification of transferrin uptake shown of images in panel (a). Scale bar is 20 µM*.

We also checked the effect of our treatment on receptor mediated endocytosis. Transferrin is a glycoprotein which plays a role in iron transport in the cells via clathrin dependent endocytosis pathway^27^. TD enters the cells via clathrin mediated endocytosis^20^. So, we decided to check the cellular uptake of transferrin-A647 into the cells. The uptake of transferrin increased with the treatment of TD and decreased with the LPS treatment (**Figure 3c, d**). We hypothesized that transferrin uptake would increase after TD treatment in LPS treated cells. However, we found further decrease in transferrin uptake in TD + LPS and LPS + TD treated cells. The reason for this might be cell type dependent uptake of transferrin.

### 2.4 Targeting tumor associated macrophages

With establishing the role of TD in phagocytosis and the anti-oxidative and anti-inflammatory properties, we decided to explore the effect of TD in TAMs. We treated the RAW264.7 cells with conditioned medium from HeLa cells^28^. The cytokines and chemokines released from the cancerous cells will polarize the RAW264.7 cells to act like TAM. We then treated the TAM with TD to check its effect on ROS levels. We found that TAM had significantly higher levels of ROS compared to non-treated control (**Figure 4a, b**). The treatment with TD was successful in reducing the levels of ROS. We also checked for the HIFα levels in TAM and TD treated TAM. The expression pattern of HIFα was similar to the levels of ROS (**Supplementary Figure 3**). This suggests that TD can actually be used for effective treatment of TAM in tumor microenvironment.

**Figure 4:**
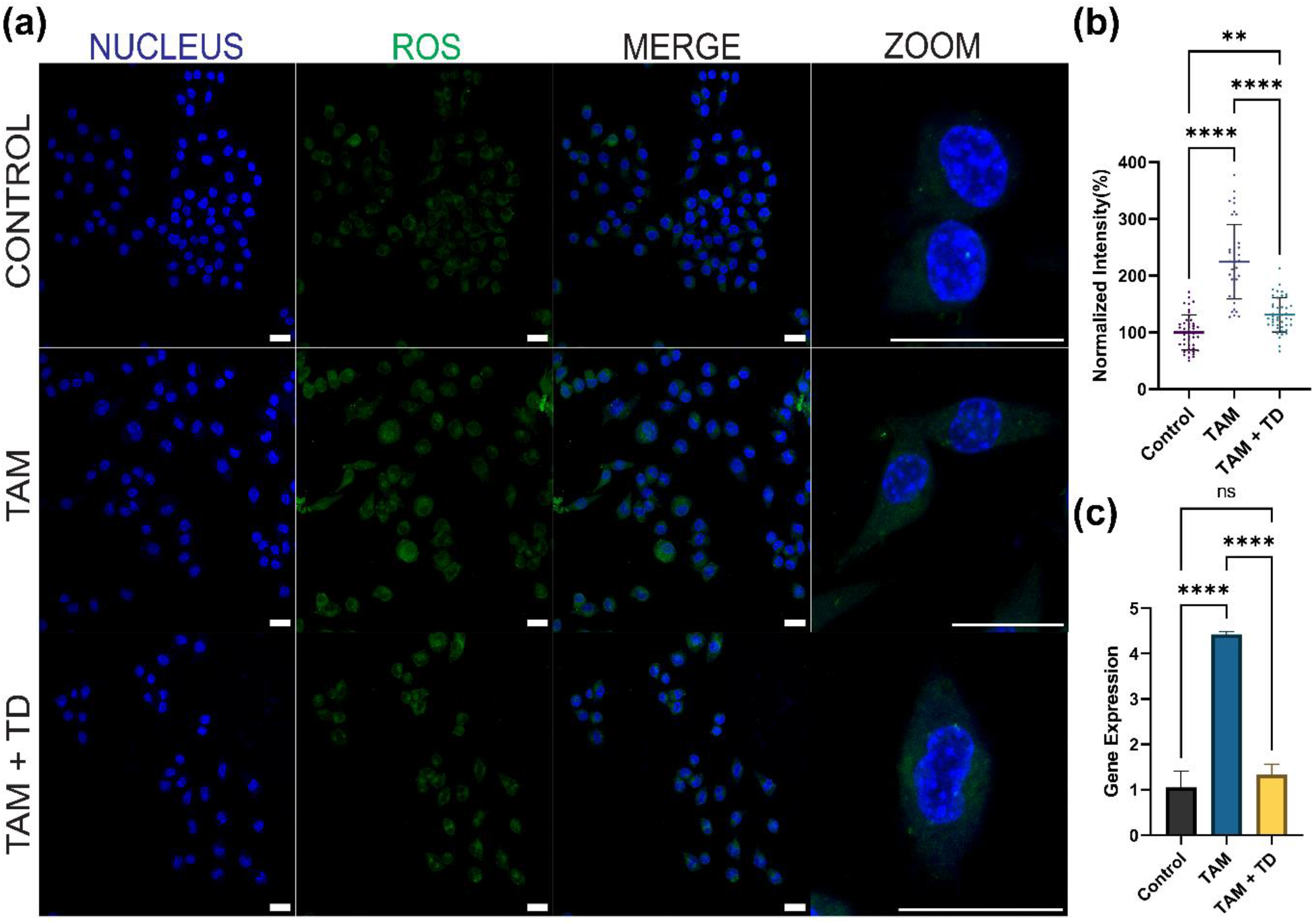
Targeting tumor associated macrophages. **(a)** Confocal images showing ROS levels using DCF-DA staining. Blue color indicates DAPI staining of nucleus, green color indicates ROS levels. **(b)** Quantification of ROS level shown of images in panel (a). **(c)** NFkB expression levels using real time PCR. Scale bar is 20 µM.

To further validate our findings, we studied the gene expression of NFkB. NFkB is shown to maintain the tumor promoting phenotype of TAM in certain cancers^29^. NFkB has been also linked to evolution of inflammatory tumors^30^. In our study, we found that in TAMs, the NFkB was highly upregulated and after TD treatment, NFkB downregulated significantly (**Figure 4c**). This confirmed our initial findings of TD as a treatment modality for TAMs.

### 2.5 TD targets PDL1 for depolarizing TAMs

To analyze the underlying mechanism for TD to target TAMs, we decided to take one step further. PD-L1 is found in healthy cells which sends signal to immune cells, particularly T cells, to not attack them^31^. Cancer cells have high amount of PD-L1, helping them to escape immune attack^32^. Multiple cytokines released by TAM can also upregulate the expression of PD-L1 by activation of JAK/STAT pathway^33^. HIF can also bind to hypoxia response element of the PD-L1 promoter region and induce the transcription of PD-L1^34^. So, we proceeded with immunostaining of PD-L1 and found the expression of PD-L1 significantly higher in TAM compared to control (**Figure 5a,b**). When treated with TD, the expression of PD-L1 decreased significantly. We also know the cell shape indicates the polarity of macrophages^35^. After TD treatment, the cell area of TAM is decreased (**Figure 5c**). This suggests that TD is depolarizing TAMs by downregulating the expression of PD-L1. Efficacy of PD-1 inhibitors are low and modulating the expression of PD-L1 might increase their efficacies.

**Figure 5:**
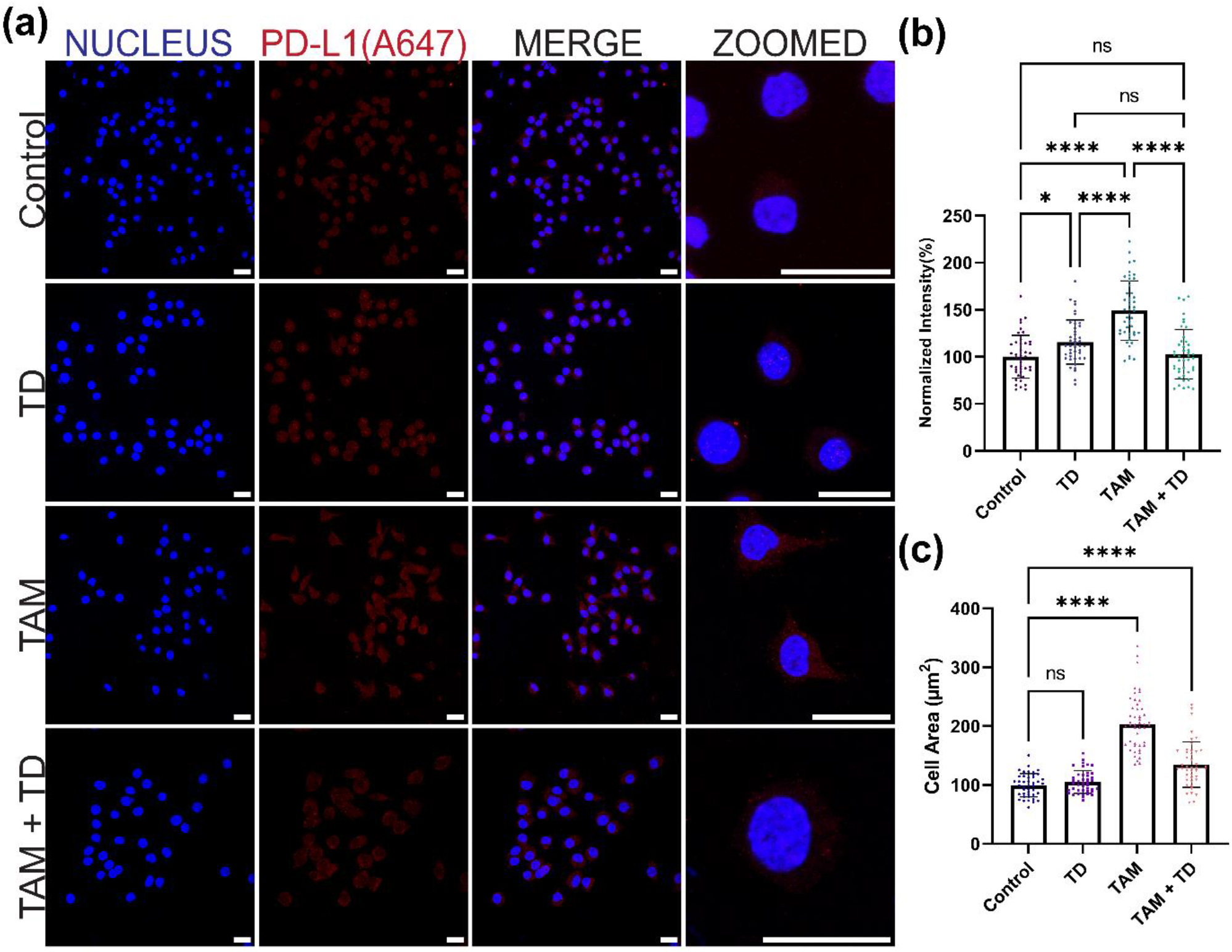
TD targets PD-L1. **(a)** Confocal images showing PD-L1 immunostaining. Blue color indicates DAPI staining of nucleus, red color indicates PD-L1 staining. **(b)** Quantification of PD-L1 level shown of images in panel (a). **(c)** Cell area of cells shown in panel (a). Scale bar is 20 µM.

## 3. Materials and Methods

### 3.1. Materials

The primers (M1-M4) were obtained from Sigma Aldrich. Nuclease free water and magnesium chloride is obtained from SRL, India. Acrylamide:Bisacrylamide (30%), TEMED, APS, Penstrap, paraformaldehyde, ethidium bromide, triton X is obtained from Himedia. 6X loading dye, 25 bp ladder, Mowiol, IL6 ELISA kit, lipopolysaccharides (LPS), DCF-DA, DAPI is obtained from Sigma Aldrich. Goat anti-rabbit A647 secondary antibody was obtained from Invitrogen. Anti-mouse IgG (H+L) F(ab)2 Fragment (A647), HIFα antibody was obtained from CST. PD L1 antibody was obtained from eBioscience. DMEM Media, FBS, trypsin was obtained from Gibco. RNA extraction kit and reverse transcription kit was obtained from Qiagen.

### 3.2. Synthesis of Tetrahedron

DNA tetrahedron was synthesized as described previously. Briefly, four single stranded oligonucleotides (Supplementary Table 1) were taken at equimolar ratio with 2 mM MgCl_2_ and denatured at 95°C for 30 minutes. They were annealed by gradually decreasing the temperature by 4°C up to 4°C for 15 minutes at each step. The final concentration of DNA tetrahedron was 2.5 µM.

### 3.3. Characterization of Tetrahedron

#### 3.3.1. Electrophoretic mobility shift assay (EMSA)

EMSA was performed for confirmation of higher order structure formation. Four tubes were taken each containing equimolar ratio of M1, M1 + M2, M1+ M2 + M3, M1+ M2 + M3 + M4 respectively. They were synthesized using the same protocol and then subjected to 8% native page at 70 V for 90 minutes. The bands were stained using ethidium bromide. The gel was visualized using Gel Documentation system (Biorad ChemiDoc MP Imaging System).

#### 3.3.2. Dynamic Light Scattering (DLS)

The size-based characterization was done using DLS. The sample was prepared by diluting TD 20-fold. Then it was subjected to Malvern analytical Zetasizer Nano ZS instrument and hydrodynamic size was measured.

#### 3.3.3. Atomic Force Microscopy (AFM)

The morphology-based characterization was performed using Bio AFM (Bruker JPK NanoWizard sense+). The sample was prepared in 1 :10 dilution on freshly cleaved mica sheet and allowed to dry. Then tapping mode was used to observe the DNA tetrahedron. The image was further processed using JPK software.

#### 3.3.4. Stability Assay

Stability of TD was checked using serum stability assay. TD was incubated with 10% FBS at 37°C for different time points (0, 1, 2, 6 hours). The reaction was stopped by immediately storing the tube at -20°C. 8% native PAGE was run to check the band intensity at 70 V for 90 minutes. The gel was visualized using Gel Documentation system (Biorad ChemiDoc MP Imaging System).

### 3.4. Cell Culture

RAW264.7 cells were kindly gifted by Dr. Ashutosh Kumar. RAW264.7 and HeLa cells were maintained in DMEM medium supplemented with 10% FBS and 1% pen-strap. They were maintained at 37°C in a humidified incubator and 5% CO_2_.

#### 3.4.1 Conditioned Media

HeLa cells were grown to 90% confluency in DMEM complete media. The medium was then collected and centrifuged at 200g for 5 minutes. The supernatant was collected and stored in -20°C until further use.

#### 3.4.2. Treatment Groups

For the first part of study, there were 5 treatment groups; control, 200 nM TD for 2 hours, 500 ng/mL LPS for 24 hours, 200 nM TD followed by 500 ng/mL LPS for 24 hours, and 500 ng/mL LPS for 24 hours followed by 200 nM TD for 2 hours. The second part of study includes three treatment groups; control, TAM (treated with conditioned media for 24 hours), TAM (treated with conditioned media for 24 hours) followed by treatment with 200 nM TD for 2 hours.

### 3.5. Reactive Oxygen Species (ROS) Detection Assay by Confocal Microscope

ROS was measured using DCF-DA staining. The cells were seeded in 24 well plate on coverslips and allowed to grow till 80% confluency followed by the treatment. The cells were washed with 1X PBS two times. They were treated with 10 µM DCF-DA for 30 minutes at 37°C and washed with 1X PBS two times. They were fixed with 4% PFA for 15 minutes at 37°C, washed three times with 1X PBS and later mounted with DAPI + Mowiol. The slides were stored at 37°C until imaging.

### 3.6. ROS using microplate reader

ROS was measured using DCF-DA treatment. The cells were seeded in 96 well plate and allowed to grow till 80% confluency followed by the treatment. The cells were washed with 1X PBS two times. They were treated with 10 µM DCF-DA for 30 minutes at 37°C and washed with 1X PBS two times. The 96 well plate was immediately subjected to microplate reader (Biotek) and reading was taken at excitation 485 nm and emission at 535 nm.

### 3.7. Immunostaining

The cells were seeded in 24 well plate on coverslips and allowed to grow till 80% confluency followed by the treatment. They were fixed with 4% PFA for 15 minutes at 37°C and washed three times with 1x PBS. They were permeabilized with 0.1% Triton-X 100 for 15 minutes at 37°C and then blocked with blocking buffer (10% FBS + 0.05% Triton-X 100) for 1 hour at 37°C. They were subjected to primary antibody at 1:100 dilution for 2 hours at 37°C followed by secondary antibody treatment for 2 hours at 37°C. The cells were then washed and mounted with DAPI + Mowiol. The slides were stored at 37°C until imaging.

### 3.8. Cellular Uptake

Cellular uptake of transferrin and dextran 40 kDa was conducted. The cells were seeded in 24 well plate on coverslips and allowed to grow till 80% confluency followed by the treatment. After the treatment, cells were incubated with transferrin 5 µg/mL or 5 mM of 40 kDa dextran for 15 minutes. The cells were washed with 1X PBS two times. They were fixed with 4% PFA for 15 minutes at 37°C, washed three times with 1X PBS and later mounted with DAPI + Mowiol. The slides were stored at 37°C until imaging.

### 3.9. ELISA

IL6 levels were measured using IL6 detection kit (Sigma). The cells were seeded in 6 well plates and treated accordingly. The supernatant was collected and centrifuged to remove any debris and then further used according to the manufacturers protocol.

### 3.10. Confocal Microscopy

Leica Sp8 confocal microscope was used for all the imaging. The slides were imaged using 63X oil immersion objective lens. The pinhole was kept at 1 airy unit. 4 lasers were used to excite different fluorophores; DAPI: 405nm, Tf-A488, DCFDA, FITC Dextran 40 kDa: 488nm, and TDCy5, anti-mouse secondary antibody A647, anti-rabbit secondary antibody A647: 633nm. 4-5 z-stacks were taken for each sample. Image analysis was done using Fiji ImageJ (NIH). The background from each image was subtracted and whole cell intensity was quantified using maximum intensity projection. Minimum 30 cells were quantified to study the cellular experiments.

### 3.11. Quantitative Real time PCR

Total RNA was isolated using Qiagen RNA isolation kit. It was reverse transcribed to cDNA using Qiagen RT-PCR kit. Target cDNA was amplified using SYBR chemistry (Applied Biosystem) in Applied Biosystems 7500 system according to the following steps: 55°C for 2 minutes, 95°C for 10 minutes, 40 cycles of (95°C for 30 seconds, 51°C for 1 minutes and 72°C for 1 minute). Melting curve was checked to detect the presence of primer dimers and false priming.

### 3.12. Statistical Analysis

All the experiments were carried out in triplicates. The data is presented as mean ± SD. One way ANOVA was carried out with Tukey’s correction using GraphPad Prism 9.0. The statistical significance is denoted by, ^*^ indicates p ≤ 0.05, ^**^ indicates p ≤ 0.01, ^***^ indicates p ≤ 0.001, ^****^ indicates p ≤ 0.0001, and ns indicates non-significant.

## 4. Conclusions

Our work focused on exploring the capabilities of TD in aspect of immunology. We found that TD is anti-oxidative and anti-inflammatory and we further explored its role in tumor associated macrophages. As per our best knowledge, this is the first work exploring the role of TD in tumor associated macrophages. We found that TD can indeed target the tumor associated macrophages by down regulating the expression of PD-L1. This work will open up the uncharted territory of using TD for not just cancer but also auto-immune diseases. TD treatment can also be explored as preventative medicine. Incorporating TD into cancer immunotherapy has a promising frontier. By targeting TAM, there is a potential to enhance the current therapeutic approaches and pave the way for new treatment modalities.

## Acknowledgements

All authors thank IITGN and CIF-IITGN for facilities. PV thanks UGC for PhD fellowship and IITGN for additional fellowship. DB thanks SERB-DST GoI and MoES-STARS for research grant, Gujcost and GSBTM for research funding. PV and DB thank Dr. Ashutosh Kumar from Ahmedabad University for kindly providing with RAW264.7 cell line.

