## Supporting information for "Designer DNA nanocages modulate anti-oxidative and anti-inflammatory responses in tumor associated macrophages"

### Supplementary Information

Payal Vaswani and Dhiraj Bhatia

Department of Biological Sciences and Engineering, Indian Institute of Technology Gandhinagar, Palaj, Gujarat, India 382355

**Table 1**: Primer sequence used for synthesis of DNA tetrahedron

| Name | Sequence (5' to 3') |
|---|---|
| M1 | ACATTCCTAAGTCTGAAACATTACAGCTTGCTACACGAGAAGAGCCGCCATA GTA |
| M2 | TATCACCAGGCAGTTGACAGTGTAGCAAGCTGTAATAGATGCGAGGGTCCA ATAC |
| M3 | TCAACTGCCTGGTGATAAAACGACACTACGTGGGAATCTACTATGGCGGCTC TTC |
| M4/M 4Cy5 | Cy5/TTCAGACTTAGGAATGTGCTTCCCACGTAGTGTCGTTTGTATTGGACCC TCGCAT |

**Table 2**: Primer sequence for real time PCR

| Name | Sequence (5' to 3') |
|---|---|
| GAPDH forward | TGCTGAGTATGTCGTGGAGT |
| GAPDH reverse | GTTCACACCCATCACAAACA |
| NFkB forward | ACAACTATGAGATGAACTCCGGG |
| NFkB reverse | CCGTGGGGCATTTTGTTCAG |

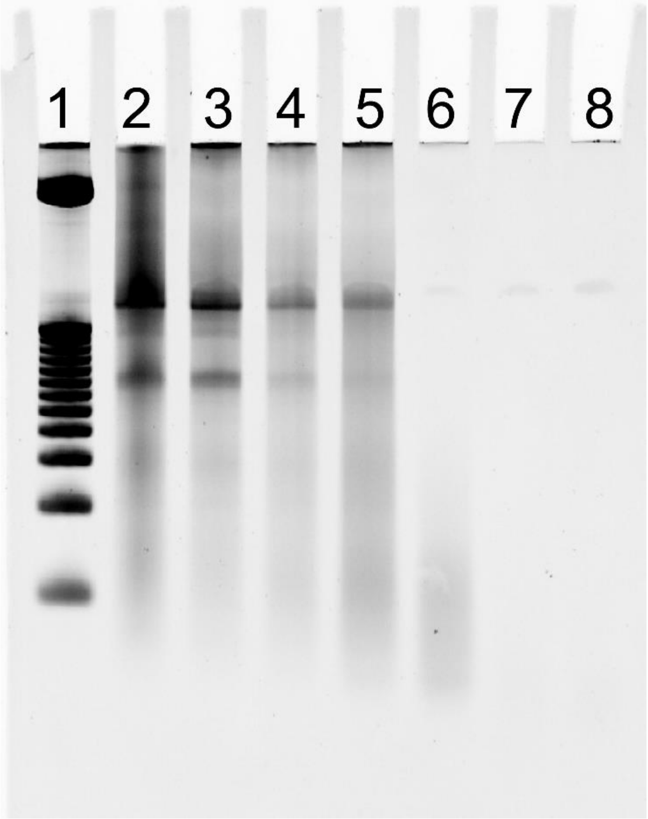


**Supplementary Figure 1: Stability of TD.** 8% native PAGE showing the stability of TD with fetal bovine serum. Lane 1: 25 bp DNA ladder; Lane 2: TD; Lane 3: 0 hour TD + 10% FBS; Lane 4: 1 hour TD + 10% FBS; Lane 5: 2 hour TD + 10% FBS; Lane 6: 6 hour TD + 10% FBS; Lane 7: 12 hour TD + 10% FBS; Lane 8 24 hour TD + 10% FBS.

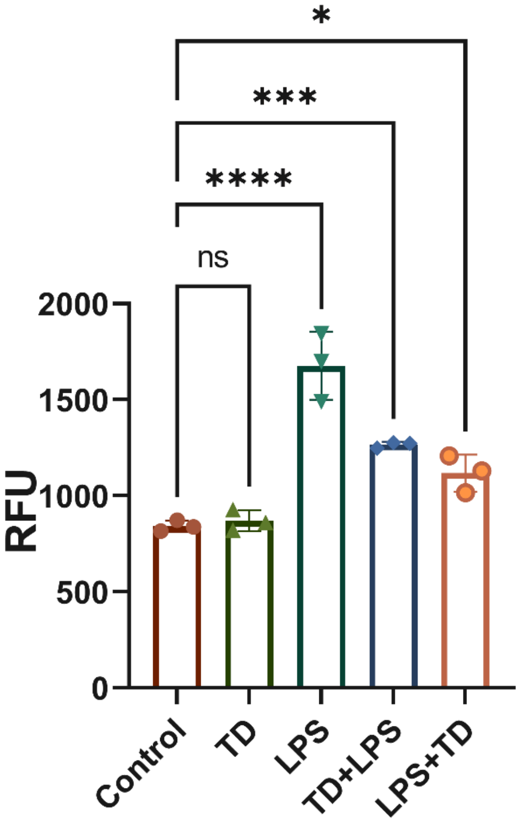


**Supplementary Figure 2: ROS levels using plate reader.** The 5 treatment groups were stained with DCF-DA dye for checking the levels of ROS. The excitation and emission were 485 nm and 535 nm respectively.

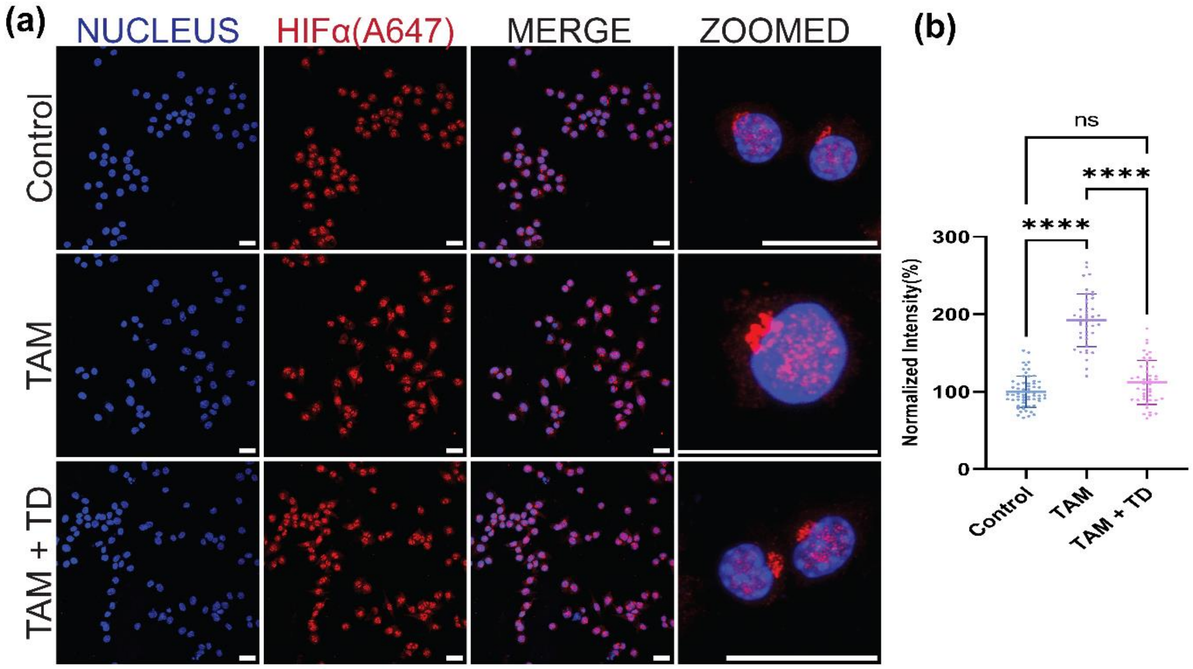


**Supplementary Figure 3: HIF-alpha immunostaining for TAM.** The RAW264.7 cells were treated with conditioned media to form tumor associated macrophages (TAM). **(a)** Confocal images showing HIFα immunostaining in control, TAM and TAM + TD group. Blue color indicates DAPI staining of nucleus, red staining indicated HIFα. **(b)** Quantification of HIFα expression in the cells in panel (a). Scale bar is 20 μM.
